# Scale-free versus multi-scale: Statistical analysis of livestock mobility patterns across species

**DOI:** 10.1101/055905

**Authors:** Takuto Sakamoto, Lloyd Sanders, Nobu Inazumi

## Abstract

In quantitative studies on animal movements and foraging, there has been ongoing debate over the relevance of Lévy walk and related stochastic models to understanding mobility patterns of diverse organisms. In this study, we collected and analyzed a large number of GPS logs that tracked the movements of different livestock species in northwestern Kenya. Statistically principled analysis has only found limited evidence for the scale-free movement patterns of the Lévy walk and its variants, even though most of the tracked movements clearly show super-diffusive behavior within the relevant temporal duration. Instead, the analysis has given strong support to composite exponential distributions (composite Brownian walks) as the best description of livestock movement trajectories in a wide array of parameter settings. Furthermore, this support has become overwhelming and near universal under an alternative criterion for model selection. These results illuminate the multi-scale and multi-modal nature of livestock spatial behavior. They also have broader theoretical and empirical implications for the related literature.

## Introduction

Over the past two decades, quantitative studies on animal movements and foraging have expanded considerably. Apart from the huge advancement in tracking technologies, one of the driving forces behind this expansion has been the ongoing debate over Lévy flight/walk and related stochastic models^1–3^. A Lévy walk is a class of random walk where a walker’s step length at each move is drawn from a so-called “fat-tailed” power law distribution (the second moment does not exist) with a certain range of exponent^4^. Ever since the seminal, somewhat controversial works by Viswanathan and his colleagues^5, 6^, arguments and counter-arguments concerning the ability of this model to describe the mobility patterns of moving organisms have prompted a great variety of theoretical^7–13^, methodological^14–17^, and empirical works^18–20^.

In this debate, a wide array of organisms has been reported to follow a Lévy walk or one of its variants (e.g., ‘truncated’ Lévy walk). Examples include cells^21^, bacteria^22^, mussels^23^, insects^6, 24^, marine predators^18, 19^, herbivores^6^, primates^25, 26^, and even ancient extinct species^27^. Human beings are no exception^2, 3, 28^. Lévy walk models have been considered to represent the movements of such diverse populations as African hunter-gatherers^29, 30^, Peruvian anchovy fishermen^31^, US travellers^32^, and UK burglars^33^. Although strong counter-examples always exist^14, 15^, mounting evidence seems to suggest that diverse living organisms actually adopt Lévy walk-like, scale-free mobility patterns, especially in harsh environments with few resources^6, 18^.

In the study reported here, we examined the applicability of these patterns to yet another group of organisms: livestock animals such as cattle, goats, sheep, and camels. These domesticated herbivores have had close relationships with human beings for thousands of years. Naturally, one might expect some characteristic traits that carry a trace of this long history in the livestock movements. Moreover, livestock animals are major contributors to human livelihood as well as its disruption, for example, by carrying deadly diseases such as sleeping sickness^34^. Thus, seeking formal descriptions of livestock mobility has practical implications not only for the animals themselves but also for human beings.

In the recent literature on animal husbandry and pastoralism, livestock movements have been actively tracked with GPS devices^35–43^. Most of these studies, however, have not been done from the dynamic perspective that pursues direct representations of the spatio-temporal behavior of livestock animals. Rather, obtained GPS data points are quickly aggregated into static summary variables such as a total walked distance and an average speed for use in ensuing statistical analyses. Furthermore, one experimental study that explicitly investigated Lévy walks in the context of goat movements^44^ suffers from a common methodological flaw that affects many other works. That is, in fitting a power law to mobility data, the authors applied linear regression to the log-transformed distribution of step lengths. Such an operation, which tends to cause a systematic bias in an estimated parameter, has largely been discredited^14, 17, 45^. This casts a serious doubt on the assumed presence of Lévy patterns that the authors derived from their analyses.

Thus, a significant gap still exists regarding the relevance of the Lévy walk or any other stochastic representations to livestock mobility patterns. We aim to bridge this gap by analyzing a large dataset on livestock movements in a statistically principled manner.

Specifically, we recorded, with GPS loggers, the movement trajectories of different livestock species during daily herding in northwest Kenya. For each of these trajectories (194 in total), we first computed diffusion statistics to see whether there could be any sign of anomalous diffusion: the first litmus test for the Lévy walk. We then converted each livestock trajectory into a series of discrete movement ‘steps’ (Fig. 1). In doing so, we followed a preceding study^30^, parameterizing various aspects of mobility in order to obtain robust results, namely, (1) a minimum distance (’minimum move’ Δ*x_min_*) between two geographic locations on a trajectory which is required for the line segment between these two locations to become a distinct step; (2) a minimum angle (’minimum turn’ Δ*θ_min_*) between two consecutive geodesic lines which is required for these two lines to become separate steps; and (3) a minimum speed (*v_min_*) of an animal between two geographic locations below which the animal is considered to pause, thus separating two movement steps. We derived distributions of movement step lengths in different combinations of these parameters (totalling 240 sets – see Methods section) for each of the 194 livestock trajectories. We then applied a set of model fitting and selection procedures to each of the step-length distributions thus derived. We investigated the descriptive power of the following six models of random walk regarding each empirical distribution: Lévy walk [power law; Eq. (1)], truncated Lévy walk [truncated power law; Eq. (2)], Brownian walk [exponential distribution; Eq. (3)], and three versions of composite Brownian walk [composite exponential distribution; Eq. (4)], namely, double-composite, triple-composite, and quadruple-composite Brownian walks. Following the relevant literature^14, 17, 30, 45^, we estimated model parameters with the maximum-likelihood estimation (MLE), screened the estimated models by means of a goodness-of-fit test that employs the Kolmogorov-Smirnov (KS) statistic, and selected the best-fit model according to Akaike’s information criterion (AIC).

**Figure 1:**
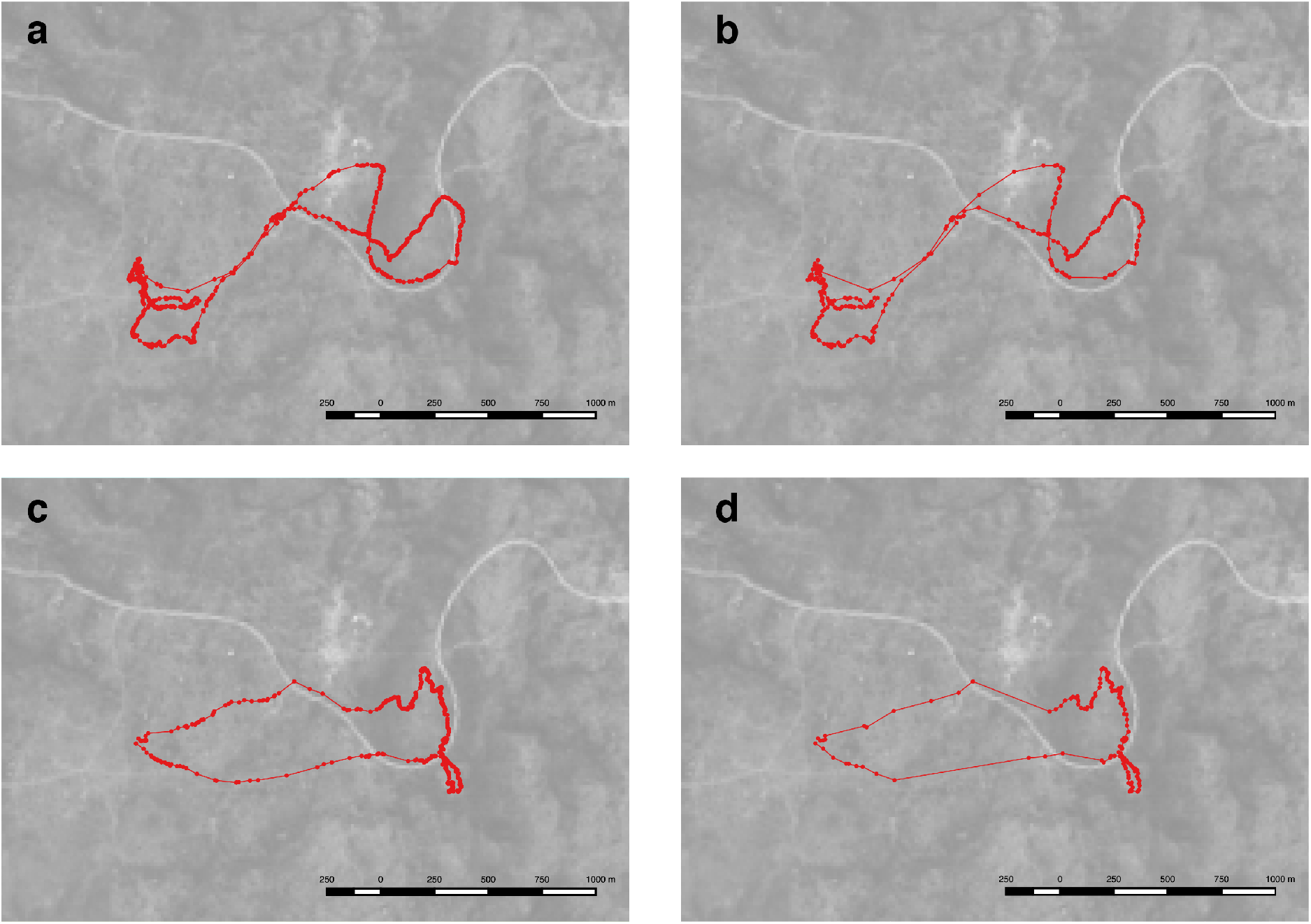
Converted livestock trajectories. The herding routes (colored red) of the livestock animals owned by one household are overlaid on a Landsat 8 satellite image of the area^46^. (**a**) Goat movement steps derived at Δ*x_min_* = 5.0m and Δ*θ_min_* = 30.0°.(**b**) A coarser version of the same trajectory obtained at Δ*x_min_* = 10.0m and Δ*θ_min_* = 60.0°. (**c** and **d**) A similar combination of images for a cattle trajectory. *v_min_* = 0.0km/h for all the cases.

## Results

### Diffusion metrics

The diffusion analysis clearly suggests the prevalence of anomalous, super-diffusive movement patterns among the recorded livestock trajectories. Figure 2 demonstrates this. Panel (a) plots the relationship between a given duration of time intervals (horizontal axis) and the mean squared displacement (MSD) of one animal which was computed for this duration (vertical axis). Panel (b) illustrates a similar plot for a different individual. These plots are log-transformed and, in each plot, a line (red) is fitted against the first 10 sample points by means of MLE. Note that incorporating data points for much longer durations can hinder accurate estimation because of the decreasing number of displacements considered in computation of MSD^47, 48^.The slope (diffusion exponent *α*) of the fitted curve then conveys important information concerning diffusion characteristics of a given trajectory: normal diffusion typified by the Brownian motion implies that *α* = 1.0; otherwise diffusion can be anomalous, either supper-diffusive (*α* ≫ 1.0) or sub-diffusive (*α* ≪ 1.0)^3^. The estimated exponents, 1.77 for (a) and 1.84 for (b), unambiguously indicate the super-diffusive regime of mobility.

**Figure 2:**
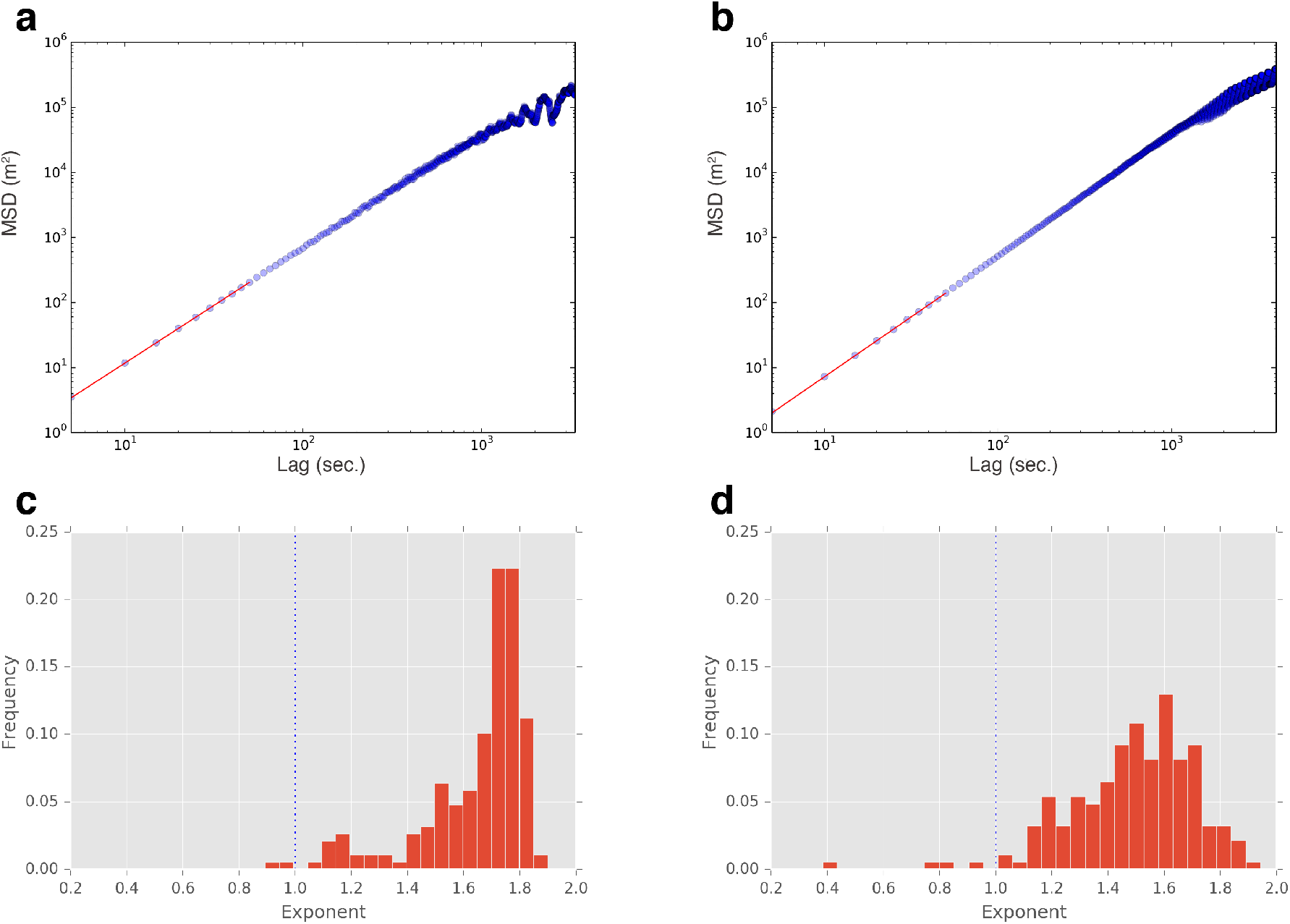
Estimated diffusion exponents show the prevalence of super-diffusion. (**a**) Log-log plots of MSDs (vertical axis) of a goat (whose trajectory is depicted in (a) and (b) of Fig. 1) against temporal lags (horizontal axis). The estimated slope of the red curve, which is fitted to the first 10 data points, is 1.77: a clear sign of the super-diffusive regime. (**b**) Similar graph for a cattle trajectory (corresponding to (c) and (d) in Fig. 1). The slope of the curve is 1.84 in this case. (**c**) Histogram of diffusion exponents in the case of 10 fitting points (sample size: 188; mean: 1.64; standard deviation: 0.20). The dotted vertical line at *α* = 1.0 indicates the boundary between the super-diffusive and sub-diffusive regimes. (**d**) Similar distribution in the case of 400 fitting points (sample size: 185; mean: 1.49; standard deviation: 0.22).

Panel (c) in Fig. 2 aggregates *α* across all the trajectories for all species in the form of a histogram. The mean of this distribution is 1.64 with the standard deviation of 0.20 (the sample size, 188, is less than the number of the recorded trajectories, 194, as some trajectories were discarded for lack of the necessary temporal duration). These numbers confirm that the above examples are not isolated outliers. The results suggest that super-diffusive movements actually comprise the dominant part of the whole data. A similar picture can be seen in Panel (d), albeit to a lesser extent. Here, the exponents were estimated using the first 400 sample points rather than 10 as in Panel (c). The mean, 1.49, is somewhat reduced from the previous one. The increased variance in *α*, denoted by the standard deviation of 0.22, partly reflects the potential increase in an estimation error noted above. Nevertheless, most of the trajectories still safely belong to the super-diffusive regime.

### Model estimation and selection

Despite the evidence for super-diffusive behavior, the model fitting and selection procedures reveal that this dominance of super-diffusive movements does not necessarily come from the frequently presumed source: the scale-free movement pattern typified by the Lévy walk. Rather, these analyses give considerable support to the composite Brownian walks against the Lévy models (the power law and the truncated power law) across livestock species. Figure 3 illustrates the overall picture (see Supplementary Fig. S1 for more complete results). Each panel in the figure is a heat map that displays relative frequencies (0.0-1.0) of a given type of model becoming the best-fit model (in the sense of minimizing AIC) for each livestock trajectory in various combinations of the mobility parameters defined in the previous section: minimum move (Δ*x_min_*) and minimum turn (Δ*θ_min_*) (see also Methods section). The remaining mobility parameter, minimum speed (*v_min_*), is fixed at 0.0 km/h [(a), (b)] or 1.0 km/h [(c), (d)]. The candidate models, whose parameters were estimated via MLE, are grouped into either composite Brownian walks [(a), (c)] or Lévy walks, including the truncated version [(b), (d)]. The simple exponential distribution is omitted because of the negligible support it received. As the figure succinctly captures, in a broad range of parameter combinations, one of the composite Brownian models (double-composite, triple-composite, and quadruple-composite versions) minimized AIC for a substantial part of step-length distributions. In particular, the descriptive power of the composite Brownian walks simply overwhelmed that of the Lévy models when *v_min_* = 1.0 km/h. Furthermore, these patterns more or less hold for each livestock species (calves, cows, and goats) separately.

**Figure 3:**
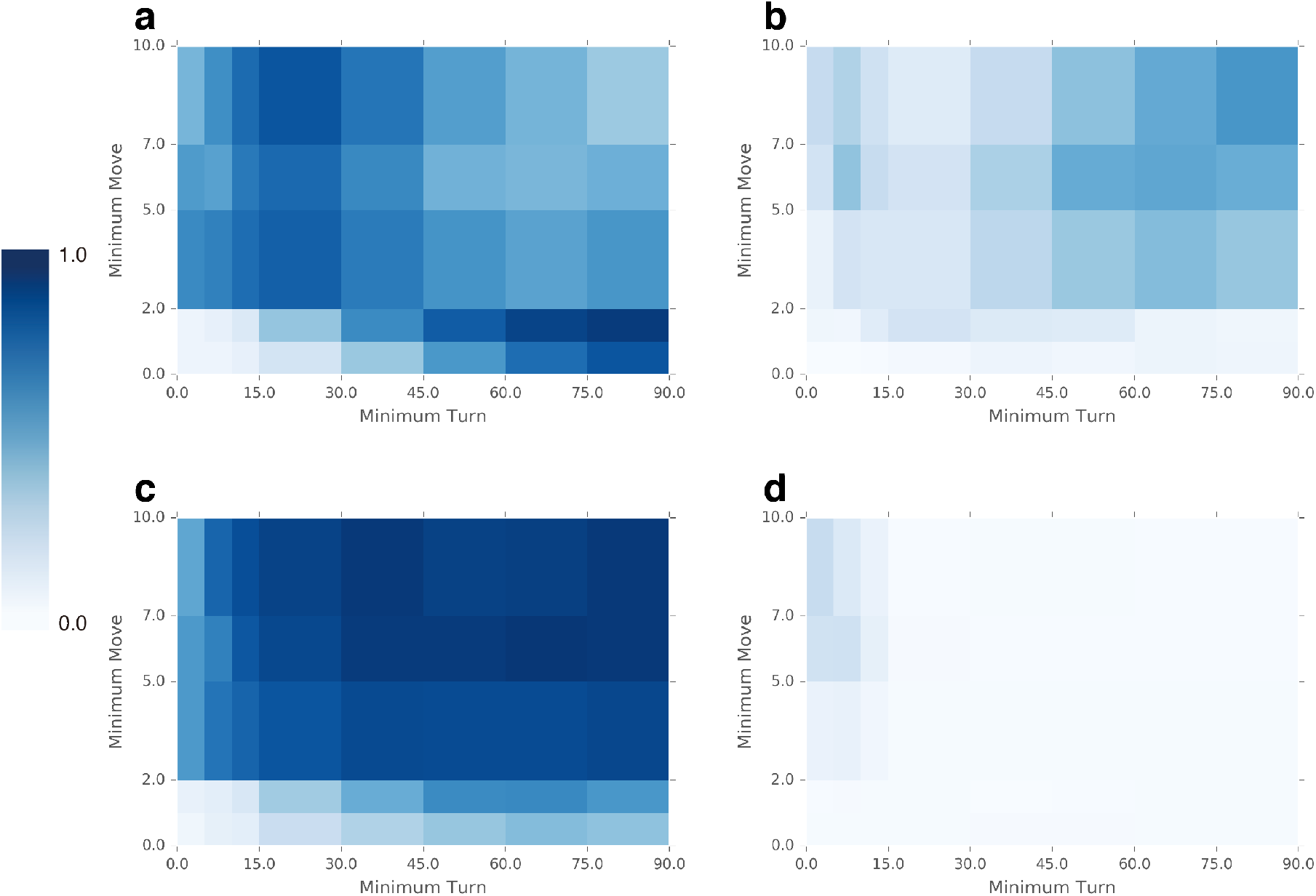
Heat maps showing best-fit frequencies. The x-axes denote changes in Δ*θ_min_*, whereas the y-axes denote changes in Δ*x_min_*. The color of each tile represents the relative frequency of one of two types of estimated random walk models, namely, composite Brownian models (double-composite, triple-composite, and quadruple-composite Brownian walks) and Lévy models (Lévy walk and truncated Lévy walk), minimizing AIC for each livestock trajectory in the corresponding combination of the parameters. (**a**) Composite Brownian models for *v_min_* = 0.0km/h. (**b**) Lévy models for *v_min_* = 0.0km/h. (**c**) Composite Brownian models for *v_min_* = 1.0km/h. (**d**) Lévy models for *v_min_* = 1.0km/h.

Figure 4 gives examples of more detailed composition of best-fit models in some combinations of Δ*x_min_* and Δ*θ_min_* for the cases of *v_min_* = 0.0km/h (see Supplementary Fig. S2 for further results). First of all, the broad ‘No Fit’ areas in Panel(a) indicate that in some conditions, where a combination of small Δ*x_min_* and small Δ*θ_min_* made derived trajectories very fine-grained and somewhat susceptible to noise, our goodness-of-fit test using the KS statistic uniformly rejected all of the six theoretical models as accurate description of most (more than 90% for Δ*θ_min_* ≤ 15.0°) of the livestock trajectories concerned. Otherwise, the relative strength of each of the models can change considerably depending on the parameters. Focusing on the comparison between the composite Brownian walks and the Lévy walks, the rough tendency is that the latter (almost exclusively the truncated version) is more likely to be selected as the best-fit model for step-length distributions that were derived with larger Δ*x_min_* and larger Δ*θ_min_*, that is, coarser data. For example, in a somewhat extreme combination of Δ*x_min_* = 10.0m and Δ*θ_min_* = 90.0°, the large majority of the derived distributions (116 out of 194, that is, 59.8%) supported the truncated Lévy walk. This most favorable situation for the Lévy models, however, lacks sufficient parallels in the other parameter combinations to overturn the overall tendency reported in the previous paragraph.

**Figure 4:**
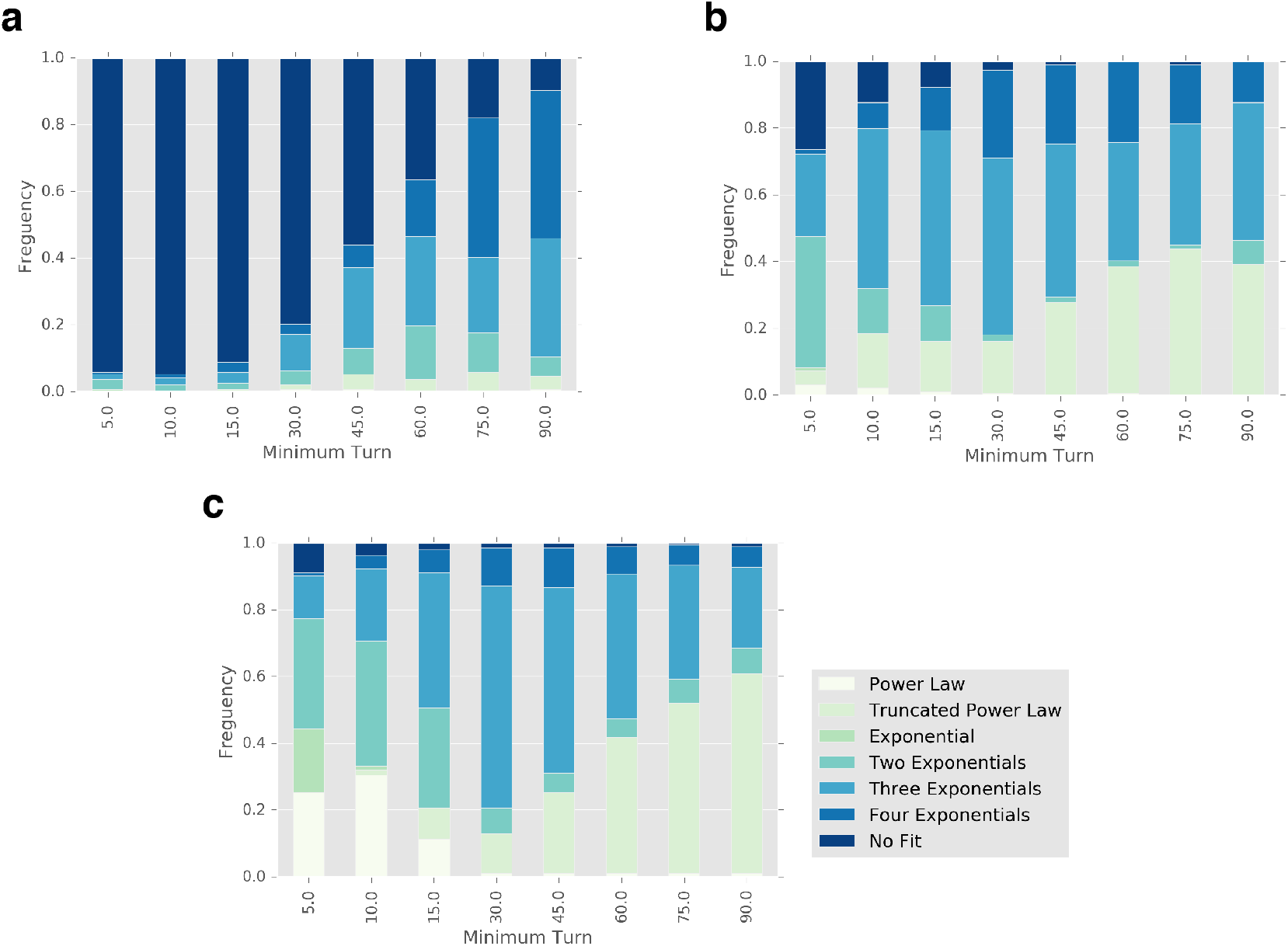
Frequency distributions of best-fit models. Each bar represents a stacked histogram (*n* = 194) that displays the frequency of each candidate model becoming the best-fit model for a given value of Δ*θ_min_*. (**a**) Δ*x_min_* = 1.0m.(b) Δ*x_min_* = 5.0m. (**c**) Δ*x_min_* = 10.0m. *v_min_* is fixed at 0.0km/h in all the histograms. ‘No Fit’ in the legend denotes the case where none of the six models passed the KS test.

Following a prominent work in the literature^30^, we also iterated the same set of analyses in the same parameter space against empirical livestock trajectories that were derived differently – namely, selecting ‘outbound’ or ‘round-trips’. Here, we reduced the dataset by redefining the trajectory of an animal as a path between its location at the start of daily herding (assumed to be 5:00 in the morning) and the furthest location from there, rather than its location at the end of herding (assumed to be 17:00 in the evening). In other words, we analyzed only the ‘outbound’ part of a trajectory instead of a ‘round-trip’ trajectory during daily herding. This change led to a considerable increase in the support the Lévy models received in some combinations of parameters, which is illustrated in Panels (a) and (b) in Fig. 5 (see also Supplementary Fig. S1). In the case of Δ*x_min_* = 5.0 m, Δ*θ_min_* = 30.0° and *v_min_* = 0.0km/h, for instance, the Lévy models were selected in 92 out of the 194 ‘outbound’ trajectories, whereas the composite Brownian models received support in 99 trajectories. The comparable figures for the ‘round-trip’ trajectories are 31 and 148, respectively.

**Figure 5:**
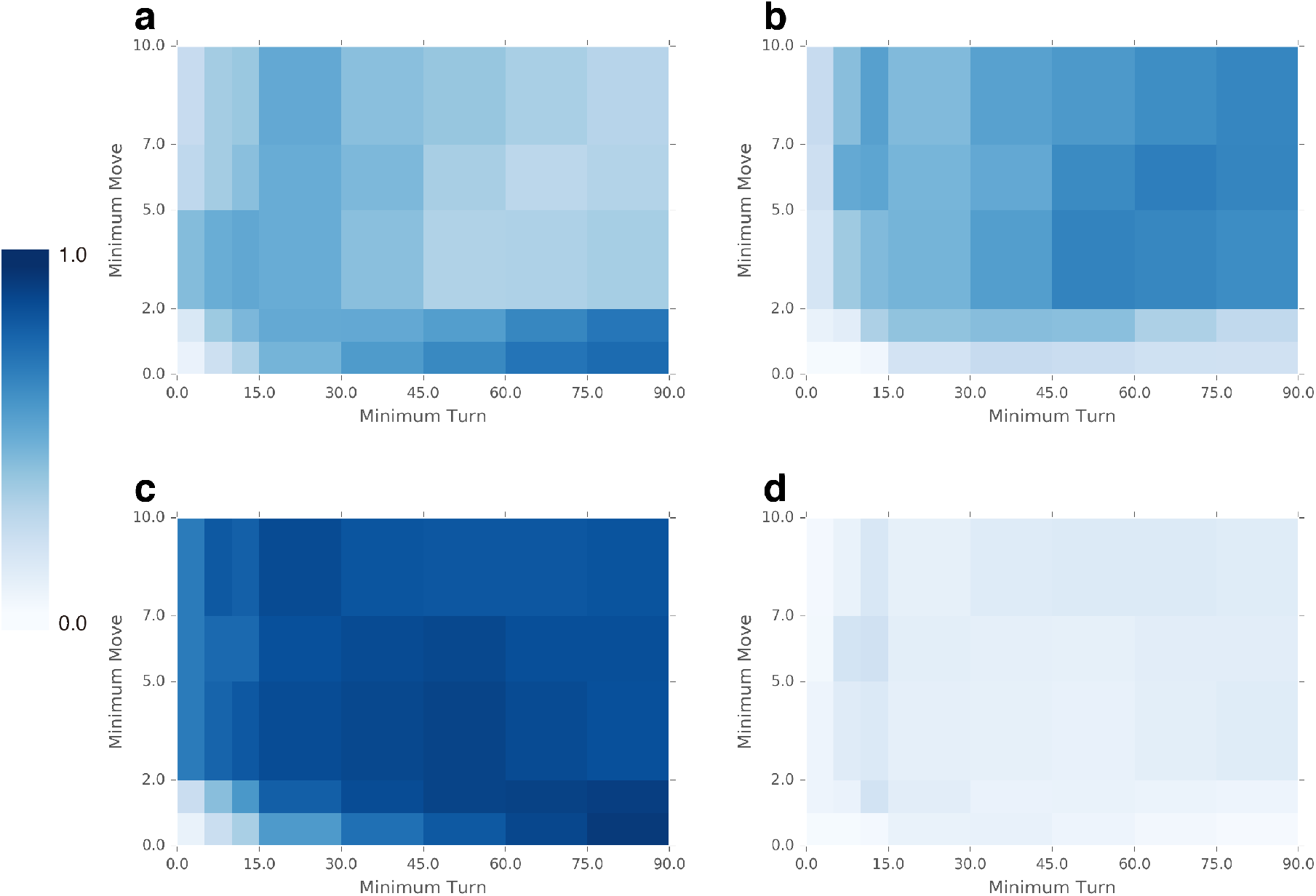
Heat maps showing best-fit frequencies in the case of ‘outbound’ trajectories. The six models were fitted against a reduced dataset where each trajectory was redefined as a path between the origin (a location at 5:00) and the furthest location from there. (**a**) Composite Brownian models for *v_min_* = 0.0km/h. (**b**) Lévy models for *v_min_* = 0.0km/h. These panels show that the support to the Lévy models significantly increases in this setting. However, the newly gained support is highly sensitive to a criterion for model selection. (**c**) and (**d**) the same as (**a**) and (**b**), respectively, except that the best-fit criterion is changed to the maximum likelihood criterion.

It should be added, however, that this increased support for the Lévy walks becomes untenable if one changes ways of perceiving the ‘best’ model. Panels (c) and (d) in Fig. 5 illuminate this point. In these graphs, we computed, in each combination of Δ*x_min_* and Δ*θ_min_*for the same ‘outbound’ dataset, the frequency in which a given type of estimated model maximized the likelihood of each step-length distribution, rather than minimized the AIC. That is, in determining best-fit models, we now focus on the accuracy of a model’s description of data regardless of the number of parameters. As the graphs clearly demonstrate, this change has more than negated the noted advantage given to the Lévy walks. The support for the composite Brownians has become overwhelming in most parameter combinations (for example, 173 out of 194 in the combination mentioned above), while that for the Lévys has been rendered almost negligible (18 in the same setting). Supplementary Figure S3 confirms that this tendency is almost universally observed in the whole parameter space, whether in the ‘outbound’ case or not.

### Estimated parameters

Among the Lévy models which were selected as the best fit models, their estimated parameters, that is, power-law exponents, typically fall within the range [*μ* < 3.0 in Eq. (2) in Methods section] where the resultant movements at least theoretically show the scale-free behavior. However, this again depends on the combination of mobility parameters employed for deriving step-length distributions. Moreover, most of the estimated exponents are substantially higher than 2.0, a ‘theoretical optimal’ in the Lévy foraging hypothesis^3, 6^, and tend towards 3.0, beyond which the movement effectively becomes single-scale Brownian motion. Panel (a) in Fig. 6 illustrates a distribution of the power-law exponents of the truncated Lévy walks selected in the case of Δ*x_min_* = 5.0 m, Δ*θ_min_* = 30.0° and *v_min_* = 0.0km/h (for the ‘round-trip’ trajectories). 30 best-fit models have their exponents around 2.72 ± 0.49 with a quarter of the exponents exceeding 3.0. A more extreme case is presented in Panel (b). In the combination of Δ*x_min_* = 7.0m, Δ*θ_min_* = 10.0° and *v_min_* = 0.0km/h, the estimated Lévy models (supported by 47 step-length distributions), all of whose exponents lie above 4.0, actually represent highly constrained movement patterns.

**Figure 6:**
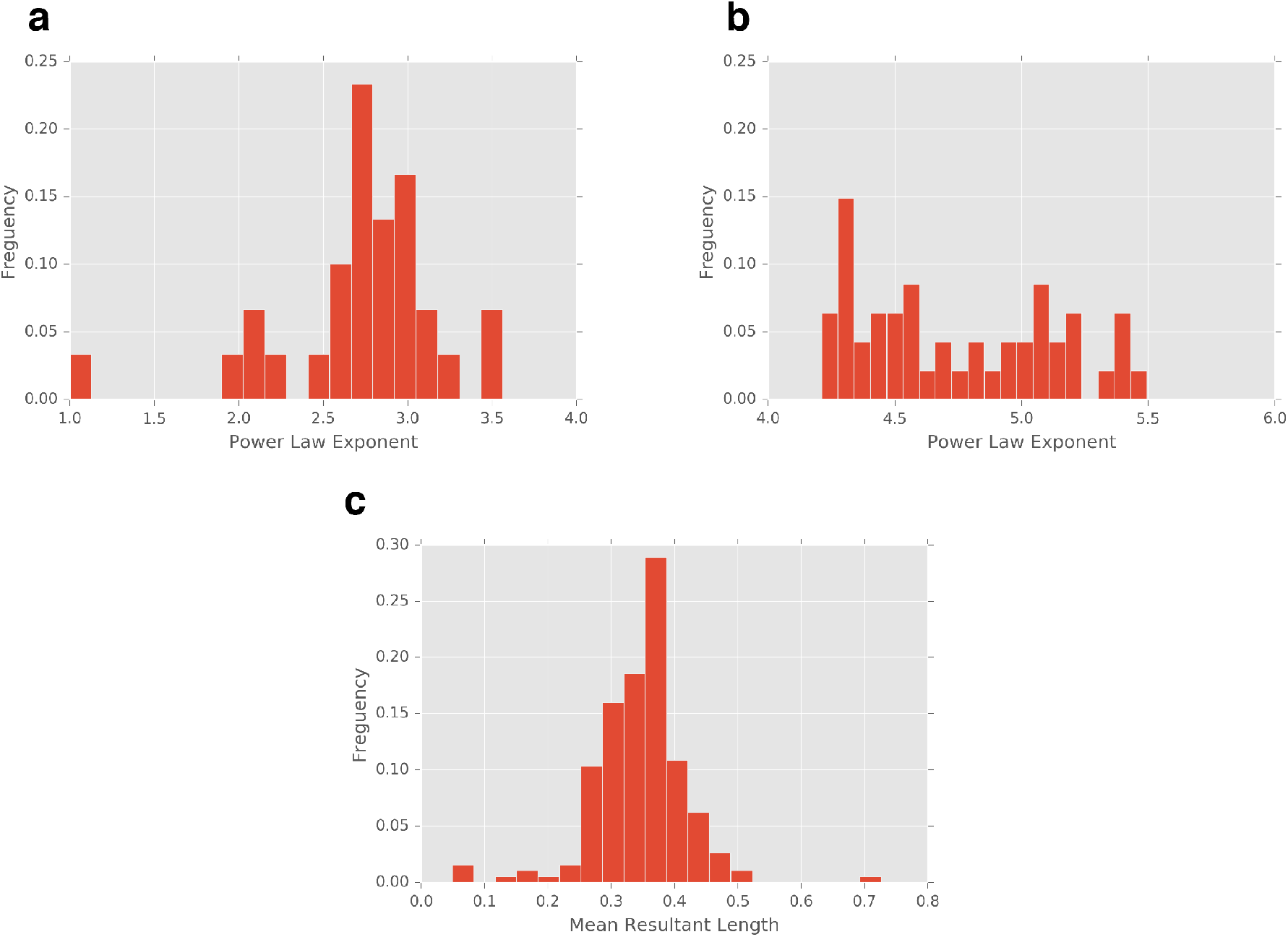
Histograms of livestock mobility metrics. (**a**) Estimated power law exponents of the truncated Lévy walk models which were selected for Δ*x_min_* = 5.0 m, Δ*θ_min_* = 30.0°, and *v_min_* = 0.0km/h (sample size: 30; mean: 2.72; standard deviation: 0.49). (**b**) Those of the Lévy walk models (non-truncated version) selected for Δ*x_min_* = 7.0m, Δ*θ_min_* = 10.0°, and *v_min_* = 0.0km/h (sample size: 47; mean: 4.75; standard deviation: 0.39). (**c**) Mean resultant lengths of turning angles which were computed for Δ*x_min_* = 5.0m, Δ*θ_min_* = 30.0°, and *v_min_* = 0.0km/h (sample size: 194; mean: 0.35; standard deviation: 0.07).

Regarding the composite Brownian models, the parameter estimation by MLE reveals the prevalence of several distinct scales of movement across livestock species in a broad array of settings. These scales of movement contribute to each step of an animal in different proportions. Supplementary Tables S1-S3 give examples of MLE parameters computed for the triple-composite Brownian model in several combinations of the mobility parameters. The tables show, for each animal, three component scales (in meters) for its move [1*/λ_j_*( *j* = 1,2,3) in Eq. (4)] (see Methods section) as well as their respective contributions [*p_j_*( *j* = 1,2,3) in the same equation]. For example, in the case of Δ*x_min_* = 5.0m, Δ*θ_min_* = 30.0° and *v_min_* = 0.0km/h, the triple-composite Brownian walk best described the majority of the step-length distributions (103 out of 194). The estimated parameters of this model (see Supplementary Table S2) indicate that the average livestock move can be decomposed into very short (0.60 ± 0.27m), moderately short (4.4 ± 1.5m), and relatively long (25 ± 16m) ranges of movement (see Discussion for further interpretation).

### Directionality of movement

Finally, we found that the movements of most livestock animals entailed a substantial degree of directional persistence: an important source of correlations in random walk which can cause highly diffusive behavior at least up to some temporal scale^3, 8, 49^. For each animal, we computed the orientation and length of the mean resultant vector from its angular turns along its trajectory^44, 50^. The former statistic denotes the circular mean of turning angles, whereas the latter (hereafter ‘mean resultant length’) measures the degree of concentration of these angles around the mean in the range of [0.0,1.0] (1.0 implies a ballistic, unidirectional motion). In other words, the mean resultant length quantifies the tendency of the animal concerned to maintain a similar heading in successive steps. Panel (c) in Fig. 6 aggregates this metric across individuals in the case of Δ*x_min_* = 5.0m, Δ*θ_min_* = 30.0°, and *v_min_* = 0.0km/h. The circular means in this case (not shown in the figure) cluster around zero (0.89 ± 13.95°). The mean resultant lengths are distributed around 0.35 ± 0.07, indicating a relatively high level of directional correlation. Similar histograms shown in Supplementary Fig. S4 confirm that this is a robust tendency observed in a broad range of parameter settings.

## Discussion

While revealing the unambiguous sign of super-diffusion in most of the livestock trajectories, we could find, at best, weak evidence for the scale-free behavior of Lévy walk as a possible mechanism for causing these diffusive movements of pastoral livestock. The Lévy models might allow a useful, parsimonious model representation of livestock mobility in some settings, for example, when analyzing a livestock trajectory on a much coarser spatial scale or focusing on some part of it such as its ‘outbound’ portion. Yet, these situations are not so predominant, in the broad range of investigations we conducted, to make any robust statement concerning the prevalence of Lévy walks. Furthermore, even within these favorable situations, the advantage given to the Lévy models, which considerably depends on the model selection criterion employed, is based on a somewhat shaky ground. Finally, the higher range of estimated power-law exponents (close to or even beyond 3.0) indicates that many of the estimated models might not be ‘strongly Lévy’ themselves.

Instead, our analyses gave broad, if not universal, support to composite Brownian walks as the best description of livestock mobility during daily herding. These models have known properties (e.g., the mixture of smaller and larger component movements) for generating, within some temporal duration, ostensibly scale-free behavior similar to that produced by the Lévy walk^10, 11^. In combination with the high directional persistence among the livestock animals revealed above, the multi-scale nature of mobility represented by these composite models seems to provide a plausible mechanism for the observed pattern of super-diffusive movement. We need, however, a more rigorous and more comprehensive analyses of empirical data to firmly establish this point.

Given the accumulation of studies that indicate the prevalence of Lévy walks in organism movements, the above results, which clearly suggest otherwise, might seem somewhat surprising. However, these results allow straightforward biological and ecological interpretations. The widespread support given to the composite Brownian walks confirms a simple fact: moving organisms have several distinctive modes of spatial behavior. Livestock animals rest, graze, walk and run during daily herding^37, 41, 51^. These animals sometimes intensively exploit nearby resources while they extensively search for distant pasture and water at other times^35^. The composite Brownian models and their estimated parameters might contain useful quantitative information on these different modes of behavior observed at multiple spatial scales.

In a much broader context, there are two ways of interpreting this apparent deviation from the assumed ubiquity of Lévy walks. Firstly, one can argue that the Lévy-type scale-free mobility pattern is not ubiquitous because many of the previous works that gave support to this pattern did so in a methodologically flawed manner. In fact, several high-profile studies on ‘Lévy foraging hypotheses’ lost ground in re-examination of their findings based on statistically enhanced approaches^5, 14, 15, 23^. These approaches, including model estimation based on MLE and model selection in a sufficiently large set of candidate models, also gave the present study a solid methodological foundation.

Moreover, we also showed that empirical support to Lévy walks could critically depend on contingent aspects of a model selection criterion. In the case of AIC, the truncated Lévy walk occasionally beat other competing models mostly because of the fewer number of parameters it has rather than the inherent accuracy of data description. The latter aspect, captured by the maximum likelihood criterion, rather gave a near-universal advantage to the composite Brownian walks. Although AIC is one of the established statistics in model selection and has also been widely employed in the preceding studies on organism movements^14, 18, 23, 30^, it is not necessarily an obvious choice. Other selection criteria such as those based on the likelihood ratio should also be tried in selecting best-fit models^45, 52^.

On the other hand, one can also claim that there is something special about domesticated animals, which might contribute to any systematic discrepancy in movement patterns between livestock and other ‘wild’ organisms. For example, livestock animals, even if extensively herded under open access or communal land tenure regimes, typically have some limited grazing ranges (‘herding radius’) set by their owners for management or other social purposes (e.g., conflict avoidance)^37, 41, 53, 54^. These ranges, normally encircling main homesteads and water points, can effectively constrain the inherent mobility of the animals concerned. Thus, possibly except in extreme circumstances such as severe drought, these animals might be unlikely to show extraordinarily long jumps in their movement trajectories, which characterize scale-free mobility patterns such as the Lévy walk. The increasing trend toward sedentarization in the contemporary pastoralism ^55^ seems to make these considerations even more important.

The two interpretations discussed so far are not mutually exclusive and both are worth exploring by further analysis. Sound methodologies combined with broad consideration of causal mechanisms can lead to balanced assessment of movement patterns of living organisms.

## Methods

### Data collection

We tracked livestock movements during daily herding in northwest Kenya from 2012 to 2014. The study area surrounds a small town named Tangulbei (0°48’0”N 36°17’12”E) in Baringo County, and 20 Pokot households residing there collaborated with the data collection. The data was collected in a part of anthropological research on pastoral livestock management, which was approved by the faculty of the Graduate School of Asian and African Area Studies (ASAFAS) of Kyoto University, with which one of the authors (Nobu Inazumi) affiliates. In conducting this study, we fully complied with relevant regulations and guidelines in anthropological field research. Most importantly, we obtained informed consent from all of the pastoral households who owned the livestock animals that we tracked.

In each collection trial, in each household, we attached a GPS logger (Holux wireless M-241) to one selected animal in a livestock corral in the early morning and released the individual back into its own herd. The device was retrieved in the same place in the evening after several (mostly one or two) days of daily herding. A notable feature of the pastoralism in the area is a highly autonomous herding style where human intervention remains minimum. During the trials, many of the herds, which typically consist of single species, moved, foraged, and eventually returned home mostly on their own, sometimes even without any human presence at all. For the 20 households, we repeated such a trial 151 times (excluding failed ones such as accidental abortion of tracking) in different periods of time with the total number of days approaching 200. Most of the trials occurred from October 2012 to February 2013, but some of the data was taken as late as in June 2014. The tracked individuals, 42 in total, consists of 20 goats, 11 cattle, 10 calves, and 1 camel. In the analyses presented here, we lumped together these different species in order to obtain an extensive sample of data. The GPS loggers recorded the locations of each individual at 5-second intervals. The total number of the data points obtained amounts to 2,126,457.

We further trimmed this large volume of track records to obtain a collection of relevant livestock trajectories. Namely, for each set of the GPS logs that were recorded in each data collection trial, we extracted its subsets according to the date and the time of measurement. Regarding the measurement time, we used only data points that were recorded during 5:00-17:00, as typical daily herding takes place within this duration. The resulting dataset consists of 194 daily livestock trajectories, containing a total of 1,565,782 data points.

### Diffusion analysis

For each of the 194 trajectories, we examined the scaling behavior of its mean squared displacements (MSDs) against temporal lags. The scale-free movement pattern of the Lévy walk implies an anomalous, super-diffusive process (but not vice versa), where MSD scales with the corresponding time lag with a power law exponent larger than unity^3^. For each possible duration of non-overlapping temporal intervals (a multiple of 5 seconds in our case), we computed the MSD of the trajectory, and, assuming the power law relationship between the lag and the MSD with Gaussian measurement noise, estimated its exponent by MLE. In this estimation, we tried different numbers of fitting points up to 800 (thus incorporating MSD computed against a maximum lag of 4,000 seconds), which is far larger than the values the preceding studies suggest as optimal (e.g., 4)^47, 48^. Though we only report statistics obtained in limited cases (10 and 400 fitting points), a particular choice in this regard does not significantly affect the diffusion characteristics of livestock movements presented here.

### Model estimation and selection

In order to make the dataset more compatible with formal representations of stochastic models, we further processed the 194 daily livestock trajectories. Specifically, for each trajectory, we discretized the location history of an individual animal into a series of movement ‘steps’. Following work on Tanzanian hunter-gatherers^30^, we define a movement step as a geodesic line segment between two locations on the recorded trajectory (1) whose length is longer than some minimum threshold value (Δ*x_min_*); (2) beyond which change in the direction of the individual exceeds some minimum threshold angle (Δ*θ_min_*); and (3) along which the speed of the individual does not fall below some minimum threshold speed (*v_min_*). Following the same work again, we also investigated the ‘outbound’ case (*δ_out_* = 1) where a movement trajectory only consists of locations between the first data point (*x*_0_) and the point farthest away from *x*_0_, in addition to the ‘round-trip’ case (*δ_out_* = 0) where all the data points are used. The four parameters [Δ*x_min_* (m), Δ*θ_min_* (°), *v_min_* (km/h), and *δ_out_*] were systematically investigated in the following ranges, generating a total of 240 possible parameter combinations.

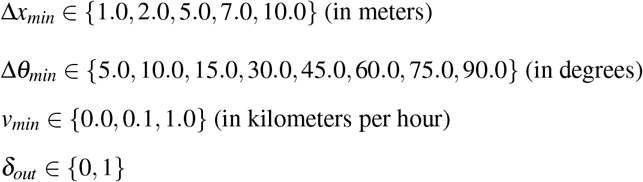

For each one of these parameter combination, we derived step-length distributions from the 194 trajectories. Employing the statistical measures developed in the preceding works^30, 45^, we then repeated model estimation and selection procedures against each of these distributions to obtain an overall distribution of best-fit models. The following random walk models, where *L* denotes the random variable for step lengths and *ℓ* its realization, were investigated as possible descriptions of livestock mobility patterns. This expanded set of models ensures a stringent assessment of the validity of a given model^15^.

Lévy walk (power law):

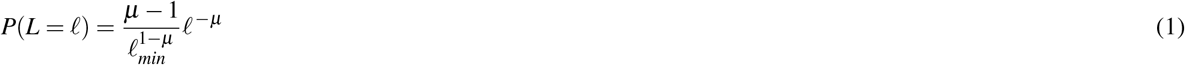

Truncated Lévy walk (truncated power law):

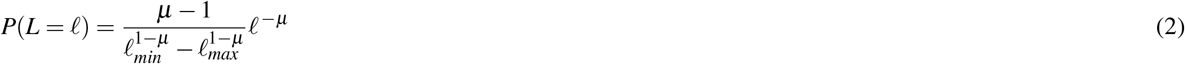

Brownian walk (exponential distribution):

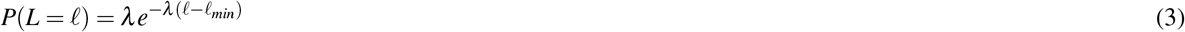

Composite Brownian walk (composite exponential distribution):

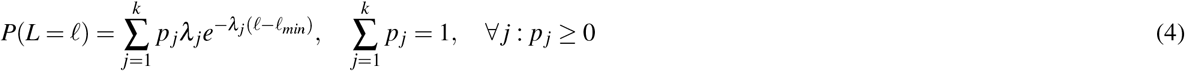

We estimated double-composite (*k* = 2), triple-composite (*k* = 3), and quadruple-composite (*k* = 4) exponential distributions.

In estimating the parameters that appear in the above equations, the left and the right truncation cutoffs (*ℓ_min_* and *ℓ_max_*) were assigned the minimum and the maximum values of a given step-length distribution, respectively^15, 30^. Regarding the other parameters, we employed the standard maximum-likelihood estimation (MLE)^14, 17, 45^. Against the random-walk models thus estimated, we further applied a two-stage model selection procedure. First, we performed a goodness-of-fit test suggested in a preceding work^45^. In this test, we quantify a divergence between a given estimated model and the corresponding empirical step-length distribution with a Kolmogorov-Smirnov (KS) statistic. We then generate, from the estimated model, a number of (500 in our case) synthetic step-length distributions with the same sample size as the empirical one, and repeat the same process of model estimation and computation of a KS statistic against each of these distributions. The original model passes the test if its KS statistic is less than the synthetic counterparts beyond a certain frequency (0.05 in our case). Second, we calculated the likelihood of the empirical step distribution under each of the remaining models and, considering the number of free-moving parameters, computed Akaike’s information criterion (AIC) for each model. The random walk model that minimizes AIC can be considered as the model that best fits the given livestock trajectory.

### Data availability

The datasets generated and analyzed during the current study are available in an anonymized form from the corresponding author on reasonable request.

## Acknowledgements (not compulsory)

The authors, N.I. in particular, wish to thank Pokot pastoralists around Tangulbei area for their generous cooperation with data collection. T.S. received financial support from the Japan Society for the Promotion of Science (JSPS) (JSPS KAKENHI; Grant 26-9525, 15KT0137, and 16K13347). The authors would also like to thank Caleb Koch and Olivia Woolley for their invaluable comments with regards to the preparation of the manuscript.

## Author contributions statement

T.S. conceived the study. N.I. collected and provided the data. T.S. and L.S. planned, coded and performed the analysis of the data. T.S. wrote the paper. All authors reviewed the manuscript.

## Additional information

The authors declare no competing financial interests.

## References

1. Pyke, G. H. Understanding movements of organisms: it’s time to abandon the Lévy foraging hypothesis.Methods Ecol. Evol.6, ^1–16^ (2015). URL http://dx.doi.org/10.1111/2041-210X.12298http://onlinelibrary.wiley.com/store/10.1111/2041-210X.12298/asset/mee312298.pdf?v=1&t=ia1n73a0&s=72c0000232f03397c03e7c040cd0741caf9431c0. DOI 10.1111/2041-210X.12298.

2. Zaburdaev, V., Denisov, S. & Klafter, J. Lévy walks. Rev. Mod. Phys.87, 483–530 (2015). URL http://link.aps.org/doi/10.1103/RevModPhys.87.483.

3. Viswanathan, G. M., da Luz, M. G. E., Raposo, E. P. & Stanley, H. E. The physics of foraging: an introduction to random searches and biological encounters(Cambridge University Press, Cambridge; New York, 2011). URL Coverimagehttp://assets.cambridge.org/97811070/06799/cover/9781107006799.jpg.

4. Tsallis, C. Lévy distributions. Phys. World10, 42–45 (1997).

5. Viswanathan, G. M. et al.Levy flight search patterns of wandering albatrosses. Nat.381, 413–415 (1996). URL http://dx.doi.org/10.1038/381413a0.

6. Viswanathan, G. M. et al.Optimizing the success of random searches. Nat.401, 911–914 (1999). URL http://dx.doi.org/10.1038/44831http://www.nature.com/nature/journal/v401/n6756/pdf/401911a0.pdf.

7. Bartumeus, F., Catalan, J., Fulco, U. L., Lyra, M. L. & Viswanathan, G. M. Optimizing the encounter rate in biological interactions: Levy versus brownian strategies. Phys Rev Lett88, 097901 (2002). URL http://www.ncbi.nlm.nih.gov/pubmed/11864054 http://journals.aps.org/prl/abstract/10.1103/PhysRevLett.88.097901. DOI 10.1103/PhysRevLett.88.097901.

8. Bartumeus, F., da Luz, M. G. E., Viswanathan, G. M. & Catalan, J. Animal search strategies: a quantitative random-walk analysis. Ecol.86, 3078–3087 (2005). URL http://dx.doi.org/10.1890/04-1806http://www.esajournals.org/doi/pdf/10.1890/04-1806. DOI 10.1890/04-1806.

9. James, A., Plank, M. J. & Brown, R. Optimizing the encounter rate in biological interactions: Ballistic versus Levy versus Brownian strategies. Phys Rev E Stat Nonlin Soft Matter Phys78, 051128 (2008). URL http://www.ncbi.nlm.nih.gov/pubmed/19113116 http://journals.aps.org/pre/abstract/10.1103/PhysRevE.78.051128. DOI 10.1103/PhysRevE.78.051128.

10. Plank, M. & James, A. Optimal foraging: Lévy pattern or process? J. The Royal Soc. Interface5, 1077–1086 (2008). URL http://rsif.royalsocietypublishing.org/royinterface/5/26/1077.full.pdfhttp://rsif.royalsocietypublishing.org/content/royinterface/5/26/1077.full.pdf. DOI 10.1098/rsif.2008.0006.

11. Benhamou, S. How many animals really do the Lévy walk? Ecol.88, 1962–1969 (2007). URL http://www.jstor.org/stable/27651327.

12. Lomholt, M. A., Tal, K., Metzler, R. & Joseph, K. Lévy strategies in intermittent search processes are advantageous. Proc. Natl. Acad. Sci.105, 11055–11059 (2008). URL http://www.pnas.org/content/105/32/11055.abstracthttp://www.pnas.org/content/105/32/11055.full.pdf. DOI 10.1073/pnas.0803117105.

13. Boyer, D. & Walsh, P. D. Modelling the mobility of living organisms in heterogeneous landscapes: does memory improve foraging success? Philos. Transactions Royal Soc. Lond. A: Math. Phys. Eng. Sci.368, 5645–5659 (2010). URL http://rsta.royalsocietypublishing.org/roypta/368/1933/5645.full.pdfhttp://rsta.royalsocietypublishing.org/content/roypta/368/1933/5645.full.pdf. DOI 10.1098/rsta.2010.0275.

14. Edwards, A.m. et al.Revisiting levy flight search patterns of wandering albatrosses, bumblebees and deer. Nat.449, 1044–1048 (2007). URL http://dx.doi.org/10.1038/nature06199.

15. Jansen, V. A. A., Mashanova, A. & Petrovskii, S. Comment on “Lévy walks evolve through interaction between movement and environmental complexity”. Sci.335, 918 (2012). URL http://www.sciencemag.org/content/335/6071/918.3.abstracthttp://classic.sciencemag.org/content/335/6071/918.3.full.pdf. DOI 10.1126/science.1215747.

16. Humphries, N. E., Weimerskirch, H. & Sims, D. W. A new approach for objective identification of turns and steps in organism movement data relevant to random walk modelling. Methods Ecol. Evol.4, 930–938 (2013). URL http://dx.doi.org/10.1111/2041-210X.12096http://onlinelibrary.wiley.com/store/10.1111/2041-210X.12096/asset/mee312096.pdf?v=1&t=ihprcp18&s=52d890e4085ea444dbd1d694b63e793933526627. DOI 10.1111/2041-210X.12096.

17. White, E. P., Enquist, B. J. & Green, J. L. On estimating the exponent of power-law frequency distributions. Ecol.89, 905–912 (2008). URL http://dx.doi.org/10.1890/07-1288.1http://onlinelibrary.wiley.com/doi/10.1890/07-1288.1/abstract. DOI 10.1890/07-1288.1.

18. Humphries, N. E. et al.Environmental context explains Lévy and brownian movement patterns of marine predators. Nat.465, 1066–1069 (2010). URL http://dx.doi.org/10.1038/nature09116http://www.nature.com/nature/journal/v465/n7301/pdf/nature09116.pdf.

19. Sims, D. W. et al.Scaling laws of marine predator search behaviour. Nat.451, 1098–1102 (2008). URL http://dx.doi.org/10.1038/nature06518http://www.nature.com/nature/journal/v451/n7182/pdf/nature06518.pdf.

20. Fauchald, P. Foraging in a hierarchical patch system. The Am. Nat.153, 603–613 (1999). URL http://www.jstor.org/stable/10.1086/303203http://www.jstor.org/stable/pdfplus/10.1086/303203.pdf?acceptTC=true. DOI 10.1086/303203.

21. Harris, T. H. et al.Generalized levy walks and the role of chemokines in migration of effector cd8+ t cells. Nat.486, 545–548 (2012). URL http://dx.doi.org/10.1038/nature11098http://www.ncbi.nlm.nih.gov/pmc/articles/PMC3387349/pdf/nihms-367690.pdf.

22. Ariel, G. et al.Swarming bacteria migrate by levy walk. Nat Commun6(2015). URL http://dx.doi.org/10.1038/ncomms9396. DOI 10.1038/ncomms9396.

23. de Jager, M., Weissing, F. J., Herman, P. M. J., Nolet, B. A. & van de Koppel, J. Lévy walks evolve through interaction between movement and environmental complexity. Sci.332, 1551–1553 (2011). URL http://science.sciencemag.org/sci/332/6037/1551.full.pdfhttp://science.sciencemag.org/content/332/6037/1551. DOI 10.1126/science.1201187.

24. Reynolds, A. M., Leprêtre, L. & Bohan, D. A. Movement patterns of tenebrio beetles demonstrate empirically that correlated-random-walks have similitude with a Lévy walk. Sci. Reports3, 3158 (2013). URL http://dx.doi.org/10.1038/srep03158. DOI 10.1038/srep03158.

25. Boyer, D. et al.Scale-free foraging by primates emerges from their interaction with a complex environment. Proc. Royal Soc. B: Biol. Sci.273, 1743–1750 (2006). URL http://www.ncbi.nlm.nih.gov/pmc/articles/PMC1634795/ http://www.ncbi.nlm.nih.gov/pmc/articles/PMC1634795/pdf/rspb20053462.pdf. DOI 10.1098/rspb.2005.3462.

26. Ramos-Fernández, G. et al.Lévy walk patterns in the foraging movements of spider monkeys (ateles geoffroyi). Behav. Ecol. Sociobiol.55, 223–230 (2004). URL http://dx.doi.org/10.1007/s00265-003-0700-6http://link.springer.com/article/10.1007%2Fs00265-003-0700-6. DOI 10.1007/s00265-003-0700-6.

27. Sims, D. W. et al.Hierarchical random walks in trace fossils and the origin of optimal search behavior. Proc. Natl. Acad. Sci.111, 11073–11078 (2014). URL http://www.pnas.org/content/111/30/11073.abstracthttp://www.pnas.org/content/111/30/11073.full.pdf. DOI 10.1073/pnas.1405966111.

28. Baronchelli, A. & Radicchi, F. Lévy flights in human behavior and cognition. Chaos, Solitons & Fractals56, 101–105 (2013). URL http://www.sciencedirect.com/science/article/pii/S0960077913001379http://ac.els-cdn.com/S0960077913001379/1-s2.0-S0960077913001379-main.pdf?_tid=2540c11a-f7db-11e4-998a-00000aab0f6b&acdnat=1431348312_eddb942af83a8f9be9edc20f47d1c348. DOI http://dx.doi.org/10.1016/j.chaos.2013.07.013.

29. Brown, C., Liebovitch, L. & Glendon, R. Lévy flights in dobe ju/’hoansi foraging patterns. Hum. Ecol.35, 129–138 (2007). URL http://dx.doi.org/10.1007/s10745-006-9083-4http://download-v2.springer.com/static/pdf/771/art%253A10.1007%252Fs10745-006-9083-4.pdf?token2=exp=1431454629~acl=%2Fstatic%2Fpdf%2F771%2Fart%25253A10.1007%25252Fs10745-006-9083-4.pdf*~hmac=d950a6941188a7a5d4081d61d68cf127d19ce2c71a18b23c9a33dd0f018965e2. DOI 10.1007/s10745-006-9083-4.

30. Raichlen, D. A. et al.Evidence of Lévy walk foraging patterns in human hunter–gatherers. Proc. Natl. Acad. Sci.111, 728–733 (2014). URL http://www.pnas.org/content/111/2/728.abstracthttp://www.pnas.org/content/111/2/728.full.pdf. DOI 10.1073/pnas.1318616111.

31. Bertrand, S., Bertrand, A., Guevara-Carrasco, R. & Gerlotto, F. Scale-invariant movements of fishermen: The same foraging strategy as natural predators. Ecol. Appl.17, 331–337 (2007). URL http://dx.doi.org/10.1890/06-0303http://www.esajournals.org/doi/pdf/10.1890/06-0303. DOI 10.1890/06-0303.

32. Brockmann, D., Hufnagel, L. & Geisel, T. The scaling laws of human travel. Nat.439, 462–465 (2006). URL http://dx.doi.org/10.1038/nature04292http://www.nature.com/nature/journal/v439/n7075/pdf/nature04292.pdf.

33. Johnson, S. D. How do offenders choose where to offend? Perspectives from animal foraging. Leg. Criminol. Psychol.19, 193–210 (2014). URL http://dx.doi.org/10.1111/lcrp.12061http://onlinelibrary.wiley.com/store/10.1111/lcrp.12061/asset/lcrp12061.pdf?v=1&t=il7mdgea&s=2316435d142fe602a2962f219130529b174bc66c. DOI 10.1111/lcrp.12061.

34. Steverding, D. The history of african trypanosomiasis. Parasites & Vectors1, 3–3 (2008). URL http://www.ncbi.nlm.nih.gov/pmc/articles/PMC2270819/ http://download.springer.com/static/pdf/908/art%253A10.1186%252F1756-3305-1-3.pdf?originUrl=http%3A%2F%2Fparasitesandvectors.biomedcentral.com%2Farticle%2F10.1186%2F1756-3305-1-3&token2=exp=1489160160~acl=%2Fstatic%2Fpdf%2F908%2Fart%25253A10.1186%25252F1756-3305-1-3.pdf*~hmac=26e3ce4404ff8122d8017b9fc203cc70baf4163857805634fe13a4531da50c4a. DOI 10.1186/1756-3305-1-3.

35. Adriansen, H. K. & Nielsen, T. T. The geography of pastoral mobility: A spatio-temporal analysis of gps data from sahelian senegal. GeoJournal64, 177–188 (2005). URL http://dx.doi.org/10.1007/s10708-005-5646-yhttp://download-v2.springer.com/static/pdf/426/art%253A10.1007%252Fs10708-005-5646-y.pdf?token2=exp=1431723919~acl=%2Fstatic%2Fpdf%2F426%2Fart%25253A10.1007%25252Fs10708-005-5646-y.pdf*~hmac=7277f83754d0ee0679134ea5b71cc4b90c8b328799bf2dbc753a15888a80b072. DOI 10.1007/s10708-005-5646-y.

36. Bertiller, M. B. & Ares, J. O. Sheep spatial grazing strategies at the arid patagonian monte, argentina. Rangel. Ecol. & Manag. 61, 38–47 (2008). URL http://www.sciencedirect.com/science/article/pii/S1550742408500170http://www.bioone.org/doi/pdf/10.2111/07-130.1. DOI http://dx.doi.org/10.2111/07-130.1.

37. Butt, B. Seasonal space-time dynamics of cattle behavior and mobility among maasai pastoralists in semi-arid Kenya.J. Arid Environ.74, 403–413 (2010). URL http://www.sciencedirect.com/science/article/pii/S0140196309003061. DOI http://dx.doi.org/10.1016/j.jaridenv.2009.09.025.

38. Butt, B. Pastoral resource access and utilization: Quantifying the spatial and temporal relationships between livestock mobility, density and biomass availability in southern Kenya. Land Degrad. & Dev.21, 520–539 (2010). URL http://dx.doi.org/10.1002/ldr.989http://onlinelibrary.wiley.com/doi/10.1002/ldr.989/abstract. DOI 10.1002/ldr.989.

39. Butt, B., Shortridge, A. & WinklerPrins, A. M. G. A. Pastoral herd management, drought coping strategies, and cattle mobility in southern Kenya. Annals Assoc. Am. Geogr.99, 309–334 (2009). URL http://dx.doi.org/10.1080/00045600802685895http://www.tandfonline.com/doi/abs/10.1080/00045600802685895. DOI 10.1080/00045600802685895.

40. Feldt, T. & Schlecht, E. Analysis of gps trajectories to assess spatio-temporal differences in grazing patterns and land use preferences of domestic livestock in southwestern Madagascar. Pastor.6, 1–17 (2016). URL http://dx.doi.org/10.1186/s13570-016-0052-2. DOI 10.1186/s13570-016-0052-2.

41. Moritz, M. et al. An integrated approach to modeling grazing pressure in pastoral systems: The case of the Logone floodplain (Cameroon). Hum. Ecol.38, 775–789 (2010). URL http://dx.doi.org/10.1007/s10745-010-9361-zhttp://link.springer.com/article/10.1007%2Fs10745-010-9361-z. DOI 10.1007/s10745-010-9361-z.

42. Raizman, E. A., Rasmussen, H. B., King, L. E., Ihwagi, F. W. & Douglas-Hamilton, I. Feasibility study on the spatial and temporal movement of samburu’s cattle and wildlife in Kenya using gps radio-tracking, remote sensing and gis. Prev Vet Med111, 76–80 (2013). URL http://www.ncbi.nlm.nih.gov/pubmed/23711505http://ac.els-cdn.com/S0167587713001451/1-s2.0-S0167587713001451-main.pdf?_tid=26469ecc-421d-11e5-8dde-00000aacb35e&acdnat=1439513046_d2fa4fd0d80abd386fa2c687025c35c1. DOI 10.1016/j.prevetmed.2013.04.007.

43. Zengeya, F. M., Murwira, A. & de Garine-Witchatitsky, M. Inference of herder presence from gps collar data of semi-free range cattle. Geocarto Int.30, 905–918 (2015). URL http://dx.doi.org/10.1080/10106049.2015.1004129http://www.tandfonline.com/doi/pdf/10.1080/10106049.2015.1004129. DOI 10.1080/10106049.2015.1004129.

44. de Knegt, H., Hengeveld, G., van Langevelde, F., de Boer, W. & Kirkman, K. Patch density determines movement patterns and foraging efficiency of large herbivores. Behav. Ecol.18, 1065–1072 (2007). URL http://beheco.oxfordjournals.org/content/18/6/1065.abstracthttp://beheco.oxfordjournals.org/content/18/6/1065.full.pdf. DOI 10.1093/beheco/arm080.

45. Clauset, A., Shalizi, C. R. & Newman, M. E. J. Power-law distributions in empirical data. SIAM Rev.51, 661–703 (2009). URL http://epubs.siam.org/doi/abs/10.1137/070710111. DOI doi:10.1137/070710111.

46. United States Geological Survey (USGS). Landsat 8 scene LC81690592013150LGN00 (2013). Obtained from USGS’s*EarthExplorer*(https://earthexplorer.usgs.gov) on November 27,2014.

47. Ernst, D. & Kohler, J. Measuring a diffusion coefficient by single-particle tracking: Statistical analysis of experimental mean squared displacement curves. Phys. Chem. Chem. Phys.15, 845–849 (2013). URL http://dx.doi.org/10.1039/C2CP43433Dhttp://pubs.rsc.org/en/Content/ArticleLanding/2013/CP/C2CP43433Dhttp://pubs.rsc.org/en/content/articlepdf/2013/cp/c2cp43433d. DOI 10.1039/C2CP43433D.

48. Michalet, X. Mean square displacement analysis of single-particle trajectories with localization error: Brownian motion in isotropic medium. Phys. review. E, Stat. nonlinear, soft matter physics82, 041914–041914 (2010). URL http://www.ncbi.nlm.nih.gov/pmc/articles/PMC3055791/https://www.ncbi.nlm.nih.gov/pmc/articles/PMC3055791/pdf/nihms273605.pdf.

49. Codling, E. A., Plank, M. J. & Benhamou, S. Random walk models in biology. J. The Royal Soc. Interface5, 813–834 (2008). URL http://rsif.royalsocietypublishing.org/royinterface/5/25/813.full.pdfhttp://d10k7sivr61qqr.cloudfront.net//content/royinterface/5/25/813.full.pdf. DOI 10.1098/rsif.2008.0014.

50. Benhamou, S. How to reliably estimate the tortuosity of an animal’s path: Straightness, sinuosity, or fractal dimension? J. Theor. Biol.229, 209–220 (2004). URL http://www.sciencedirect.com/science/article/pii/S0022519304001353http://ac.els-cdn.com/S0022519304001353/1-s2.0-S0022519304001353-main.pdf?_tid=3eb9610a-94d0-11e5-ae53-00000aacb360&acdnat=1448605963_780678954a53edfdc68bac57de362bff. DOI http://dx.doi.org/10.1016/j.jtbi.2004.03.016.

51. Schlecht, E., Hülsebusch, C., Mahler, F. & Becker, K. The use of differentially corrected global positioning system to monitor activities of cattle at pasture. Appl. Animal Behav. Sci.85, 185–202 (2004). URL http://www.appliedanimalbehaviour.com/article/S0168-1591(03)00290-9/abstract. DOI 10.1016/j.applanim.2003.11.003.

52. Vuong, Q. H. Likelihood ratio tests for model selection and non-nested hypotheses. Econom.57, 307–333 (1989). URL http://www.jstor.org/stable/1912557. DOI 10.2307/1912557.

53. Coppolillo, P. B. The landscape ecology of pastoral herding: Spatial analysis of land use and livestock production in east africa. Hum. Ecol.28, 527–560 (2000). URL http://dx.doi.org/10.1023/A%3A1026435714109http://link.springer.com/article/10.1023%2FA%3A1026435714109. DOI 10.1023/A:1026435714109.

54. Coppolillo, P. B. Central-place analysis and modeling of landscape-scale resource use in an east african agropastoral system. Landsc. Ecol.16, 205–219 (2001). URL http://dx.doi.org/10.1023/A%3A1011148503303http://link.springer.com/article/10.1023%2FA%3A1011148503303. DOI 10.1023/A:1011148503303.

55. Fratkin, E. M. & Roth, E. A. As pastoralists settle: social, health, and economic consequences of the pastoral sedentarization in Marsabit District, Kenya. Studies in human ecology (Kluwer Academic/Plenum Publishers, New York, 2004). URL Publisherdescriptionhttp://www.loc.gov/catdir/enhancements/fy0663/2004042174-d.htmlTableofcontentsonlyhttp://www.loc.gov/catdir/enhancements/fy0818/2004042174-t.html.

